# Inducible tomato defences persist in detached leaves, despite differential plant variety and gene-dependent expression

**DOI:** 10.1101/2025.09.18.677022

**Authors:** Mariya Kozak, Diogo P. Godinho, Eren Yayla, Juan Adánez, Inês Fragata, Sara Magalhães, Leonor R. Rodrigues

## Abstract

Herbivore and pathogen attacks often lead to the induction of plant defences. Given the intricate nature of such responses, plant integrity is assumed to be a pre-requisite for the successful mounting of defences, a hypothesis supported by work done in plant-pathogen interactions. However, the relevance of plant integrity in plant-herbivore interactions, particularly in direct plant defences, is unclear and lacks empirical molecular validation. To test this, we measured the expression of plant defence-related genes (PPO-D and PI-IIc, involved in the jasmonate signal pathway and PR-1a, involved in the salicylic acid signal pathway) in intact plants and detached leaves of two tomato varieties infested or not with the herbivorous spider mite *Tetranychus urticae*. We also tested whether defences persisted in detached leaves after one, four, seven or fourteen days of infestation. We found that tomato defences were induced in both intact plants and detached leaves and that the expression of all three genes in detached leaves increased over time, being the increment higher in the two late-expression genes, PI-IIc and PR-1a. Such results suggests that both intact plants and detached leaves can be used in studies addressing herbivore interactions with plant defences. However, induction levels in intact plants and detached leaves varied with plant variety and the gene assessed, ranging from higher expression in detached leaves than in intact plants to the reverse. Hence, studies aiming at identifying the role of specific genes should account for the differential, variety-dependent, role of plant integrity in their expression.

## Introduction

Plants have evolved a complex array of defences to counter herbivore attack, ranging from constitutive physical and chemical barriers (Shepherd et al. 2005; Mithöfer and Boland 2012; Pott et al. 2012) to inducible pathways, mounted only upon herbivore attack (Kant et al. 2015; Garcia et al. 2021). The latter include strategies that directly target herbivores (direct defences), such as the reinforcement of physical barriers (Traw and Bergelson 2003) and the production of toxic, repellent and anti-digestive compounds (Green and Ryan 1972; Karban and Baldwin 2007), and traits that affect herbivores indirectly (indirect defences), such as the production of volatile compounds that attract natural enemies (Heil 2008).

Inducible responses rely upon a complex signalling cascade in the plant, primary regulated by phytohormones such as jasmonic acid (JA) (Wasternack and Hause 2013), salicylic acid (SA) (Vlot et al. 2009) and ethylene (ET) (Adie et al. 2007), which are translocated from their site of origin to other parts of the plant (Atkins and Smith 2007). Hence, the induction of plant defences typically results not only in a local response at the feeding site, but it may also spread to distal undamaged plant tissues (i.e., systemic response) (Wu and Baldwin 2010; Kant et al. 2015).

Given the dynamic and complex nature of plant defence induction, plant integrity is thought to be key to the expression of local and systemic defences (Bezemer and Van Dam 2005; De Vos et al. 2005; De Vos et al. 2006; Erb et al. 2009; Van Dam and Heil 2011; Orłowska et al. 2013), namely for the transportation of the phytohormones precursors involved in the induction of defences (Gatehouse 2002; Puthoff and Smigocki 2007). Nevertheless, plants with compromised plant integrity, such as detached leaves or small leaf patches (e.g., leaflets and leaf discs), have been frequently used in plant-herbivore studies, including those addressing herbivore interactions with inducible plant defences (Dicke and Dijkman 1992; Stout et al. 1994; Koch et al. 1999; Musser et al. 2002; Sarmento et al. 2011a). Moreover, in a laboratory setting, herbivores are often maintained on detached leaves for extended periods, either during long-lasting experiments, such as experimental evolution (Fellous et al. 2014; Godinho et al. 2024) or during the maintenance of stock populations or lines (Brodeur and Cloutier 1992; Alba et al. 2015).

Some studies using fungal infections have reported significant differences between the induced response of intact plants and detached leaves, finding lower gene expression of defence genes in detached leaves than in intact plants (Wang et al. 2004; Liu et al. 2007; Orłowska et al. 2012; Orłowska et al. 2013). In contrast, studies with herbivore-plant interactions suggest that leaf detachment does not hamper the induction of plant defences (Larew 1989; Dussourd and Denno 1994; Arimura et al. 2001; Schmelz et al. 2001; Dias et al. 2022). However, out of these studies those that specifically measure gene expression only focus on indirect defences (Arimura et al. 2001; Schmelz et al. 2001), while those dealing with direct defences have only measured herbivore traits (e.g., fecundity, growth, survival), inferring the response of the plant indirectly (Larew 1989; Dussourd and Denno 1994; Dias et al. 2022). Moreover, these studies only test the short-term response of detached leaves (i.e., up to 3 days of infestation on detached leaves), neglecting the possible long-term effect of leaf senescence following excision (Smart 1994).

In the present study, we assess whether plant integrity is essential for the induction of direct plant defences after short- and long-term infestations, examining the induction of tomato plant defences (*Solanum lycopersicum*) by the phytophagous two-spotted spider mite species *Tetranychus urticae. T. urticae* is a worldwide crop pest (Attia et al. 2013) that often induces both JA and SA dependent defences of tomato plants (Alba et al. 2015) upon feeding, resulting in the production of toxic and anti-digestive compounds (Li et al. 2002; Kant et al. 2008; Rioja et al. 2017; Santamaria et al. 2020). To examine whether our findings were generalisable or idiosyncratic to the plant variety or the gene targeted, we tested the local expression of three plant defence-related genes, PPO-D and PI-IIc, induced by the JA signal pathway, and PR-1a, induced by the SA signal pathway, in two commonly used tomato varieties, Moneymaker and Castlemart, after two days of infestation of intact and detached tomato leaves. These genes have been commonly used to assess plant defence induction by herbivores in studies both using intact plants and detached leaves (Sarmento et al. 2011a; Alba et al. 2015). Additionally, to determine whether the induced local response in detached leaves is maintained for an extended period, we measured gene expression of plant defence-related genes following one, four, seven and 14 days of infestation, encompassing the whole life cycle of spider mites (10-12 days from egg to adult at 25°C).

## Material and methods

### Plant varieties, spider mite population and rearing conditions

Tomato plants (*Solanum lycopersicum*) of the varieties Moneymaker (MM, hereafter) and Castlemart (CM, hereafter) were sown in soil (SIRO) with or without fertilizer (BoskGrow Mixcote, NPK 10-10-22), respectively, following a standard protocol, and watered three times a week. Bean plants (*Phaseolus vulgaris*, variety Speedy) were also grown and used to maintain the spider-mite population. Plants were grown in an herbivore-free climatic chamber (25 ± 2°C, 60% RH, 16:8h L:D) for five weeks, in the case of tomato plants, and two weeks, in the case of bean, before being exposed to spider mites.

A population of *Tetranychus urticae (*Santpoort-2; Alba et al. 2015, originally described in Kant et al. 2008 as “KMB”) was maintained in large numbers (>1000) on intact bean plants under controlled conditions (25 ± 2°C, 60% RH, 16:8h L:D) prior to the onset of the experiments. This population is a benchmark control for the induction of plant defences (Sarmento et al. 2011b; Alba et al. 2015).

Two weeks prior to each experiment, 110 adult *T. urticae* females were placed in plastic boxes (16.5 x 16.5 x 12 cm) containing two detached bean leaves with their petioles in a pot with water. The females laid eggs for two days and were then removed using a brush to prevent plant damage. Ten days later, the adult offspring, all approximately of the same age, were used in the experiment.

### Experiments

#### a) Plant defence induction in intact plants and detached leaves

To determine whether local defence induction by spider mites is contingent upon plant integrity, the expression of defence genes on infested intact CM and MM plants was compared with that on infested detached leaves. In the latter treatment, leaves were excised with scissors and the petiole was placed in a small plastic pot containing water. In both treatments, the fourth leaf (counting from the roots up to the tip of the plant) was infested with 100 mites (20 per leaflet) and isolated using a thin barrier composed of a mixture of lanolin and insect glue (1:1) around the petiole (Fig.1). Non-infested plants and leaves were used as control treatments. Intact plants and detached leaves were maintained inside closed plastic boxes (56 x 39 x 42 cm and 16.5 x 16.5 x 12 cm, respectively) with lids that contained an opening with polyamid mesh (80 µm width) to allow airflow, under controlled conditions (25 ± 2°C, 60% RH, 16:8h L:D). After two days of infestation, the females were removed, and a nine mm Ø leaf disc was cut from the second leaflet (counting from the basis of the fourth leaf counterclockwise adaxial facing) close to the petiole (Fig.1). In the treatment with intact plants, the leaf was detached from the plant immediately before this step, such that all biochemical interactions between the mites and the plant occurred on the intact plant. The leaf disc was placed in a sterile SafeLock 2mL tube containing three glass beads and immediately immersed in liquid nitrogen. The tubes were stored at -80°C for later molecular analyses (Fig.1a). Each treatment and respective control of CM and MM plants were tested in one and two blocks, respectively, with three replicates per block (Table S1a).

**Fig. 1.**
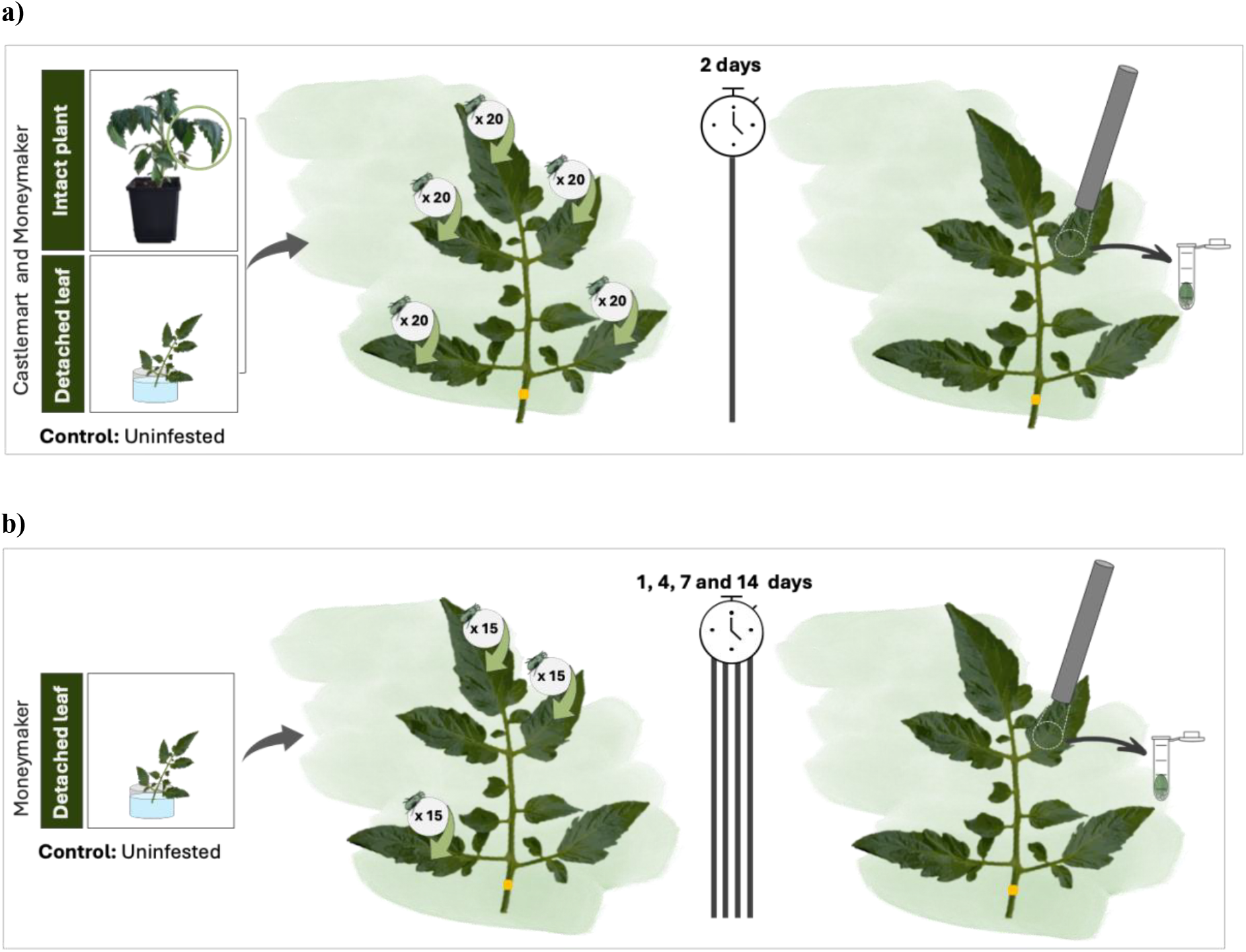
Illustration of the experiments performed to assess a) plant defence induction comparing intact plants and detached leaves and b) the persistence of defence induction in detached leaves. In a), after isolating the leaf, each leaflet was infested with 20 *T. urticae* for two days. In b), three random leaflets were infested with 15 *T. urticae* adult females for one, four, seven or 14 days. In both experiments, a nine mm Ø leaf disc was cut from the second leaflet after mite removal and immediately frozen in liquid nitrogen. The mixture of equal parts of lanolin and insect glue that was used to restrain mites from escaping is depicted in yellow around the petiole.

#### b) Long-term plant defence induction in detached leaves

To determine whether the induction of plant defences lasts for long periods after leaf detachment, the expression of defence genes was analysed after one, four, seven and 14 days of infestation of detached leaves with *T. urticae*. This experiment was performed using the third, fourth and fifth detached leaves of five-week-old MM plants, all detached on the same day and individually placed inside a small plastic pot with water. Then, forty-five adult females were randomly distributed across three leaflets of a single leaf (i.e., fifteen females per leaflet) across all time point treatments. After the established infestation period, the females were removed, and a nine mm Ø leaf disc was cut and stored as described in the previous experiment (Fig.1b). A control treatment for the absence of plant defence induction, consisting of leaves not infested with mites, was included for each time point. All leaves were kept in small plastic boxes (16.5 x 16.5 x 12 cm) under controlled conditions (25 ± 2°C, 60% RH, 16:8h L:D). For each infestation period and respective control, nine replicates (three per leaf age) were tested (Table S1b).

### RNA extraction and gene expression analysis

Total RNA was isolated using a protocol adapted from Verwoerd et al. 1989, with minor modifications (see supplementary information). After isolation, 2µg of RNA were DNAse-treated with Ambion Turbo DNA-free kit (Invitrogen) and cDNA was synthesized with RevertAid H Minus Reverse Transcriptase (Thermo Fisher Scientific). Next, 1µL of cDNA was used as template for a 20µL quantitative reserve-transcriptase polymerase chain reaction (qRT-PCR) using the CFX96 Real-Time system (Bio-Rad) with SsoFast™ EvaGreen®Supermix (Bio-Rad). The expression of three tomato defence-related genes - PPO-D, PI-IIc and PR-1 - was measured (Table S2). PPO-D and PI-IIc are marker genes of the JA pathway (Felton et al. 1989; Farmer et al. 1992), while PR-1a is a marker gene of the SA pathway (Van Loon and Van Strien 1999). The first is considered as an early gene (i.e., exhibiting maximum expression one day post-infestation), and the two latter as late genes (i.e., displaying increased expression until the leaf death) (Alba et al. 2015).

The tomato Actin gene (Table S2) was used as a reference housekeeping gene to calculate the normalized expression (NE), based on ΔCt method, as described in (Alba et al. 2015). NE values were scaled to the treatment with the lowest average NE. Observations in which the reactions failed to reach the minimum signal intensity at 40 cycles of qPCR were removed from the analysis. Three technical replicates were tested per sample.

### Statistical analyses

All statistical analyses were performed using the software R (version 4.4.1, 2024). Prior to calculating the NE, outliers were identified using the Rosner test (*rosnerTest* function from the *EnvStats* package; version 2.7.0; (Millard 2013)). Given the considerable distance between points in some treatments, the NE values were log-transformed, facilitating the visualization of the results. Results of the expression of plant defence-related genes were presented and analysed as the difference in the expression induced by *T. urticae* population to the expression in uninfested plants or leaves (i.e., the control treatment). To obtain the values of relative expression, the average NE of uninfested plants or leaves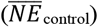was calculated for each variety within block and then the respective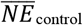 was subtracted to the NE calculated for each sample. Exceptionally, in the experiment comparing the defence induction in intact plants and detached leaves, the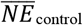 of PI-IIc gene in CM intact plants was set to zero because all reactions failed to reach the minimum signal intensity within 40 cycles of gene amplification. Graphical representations of the data were produced using the *ggplot2* package (version 3.3.6; (Wickham 2016)).

Prior to analyses comparing the expression of each gene in intact plants and detached leaves, the normality and homoscedasticity of the residuals were assessed performing Shapiro-Wilk’s (Shapiro and Wilk 1965) and Levenes’s (Levene 1960) tests. Then, linear mixed-effect models (*glmmTMB* function of the *glmmTMB* package; version 1.1.29; (Brooks et al. 2017)) were performed for each gene. The interaction between treatment (intact plant and detached leaf) and plant variety (MM and CM) were included as fixed factors in all models, while the interaction between plant variety and block was included as random factor at the intercept level (Table S3a); models with and without random factor were compared using the *anova* function (Chambers and Hastie 1992), retaining the one with the lowest Akaike Information Criterion (AIC). On that model, the significance of the interaction between fixed factors was determined using the *Anova* function (*car* package; version 3.1-2; type=III; (Fox and Weisberg 2018)). Whenever a significant interaction between fixed factors was observed, comparisons were performed between treatments within each plant variety, using the *contrast* function of the *emmeans* package (version 1.8.5; (Russell V. Lenth 2020)), with false discovery rate (FDR) corrections (Benjamini and Hochberg 1995). Finally, to test whether defence induction in plants infested with *T. urticae* differs from the control treatment (i.e., uninfested plants), we tested whether the confidence limits of the relative NE of each gene overlapped with 0 using the *test* function of the *emmeans* package.

In the analyses comparing the gene expression of detached leaves infested with *T. urticae* at different time points, the test of normality and homocedasticity of the residuals, as well as the criteria for exclusion of random factors, were applied as described above. These linear mixed-effect models included day (one, four, seven and 14) as covariate and leaf number (third, fourth and fifth) as random factor (Table S3b). To determine the trend of gene expression along time, the slope for each gene was calculated using the *summary* function (version 4.4.1; (Chambers and Hastie 1992)). Finally, the comparisons between *T. urticae* infested and uninfested treatments were performed as described above.

## Results

### Induction of plant defences occurs in both intact plants and detached leaves and differs between plant variety in a gene-dependent manner

The impact of *T. urticae* infestation on the expression of the genes studied varied with both plant treatment and variety. The relative expression of PPO-D was higher in MM than in CM plants (Fig.2a; Table S4; plant treatment x plant variety interaction: χ2=0.9585, *p-value*=0.3276; plant variety: *χ^2^*=8.3925, *p-value*=0.0038) and generally higher in intact plants leaves than in detached leaves (Fig.2a; Table S4; plant treatment: χ2=5.8941, p-value=0.0152), particularly so in MM plants (Fig.2a; detached (CM) – intact (CM): *t-ratio*=-0.639, *p-value*=0.53; detached (MM) – intact (MM): *t-ratio*=-2.600, *p-value*=0.04). The relative expression of PI-IIc, in turn, was higher in detached leaves than in intact plants, but only in CM plants (Table S4; plant treatment x plant variety interaction: χ2=5.9013, *p-value*=0.0151; Fig.2b; CM plants: *t-ratio*=3.592, *p-value*=0.0066; MM plants: *t-ratio*=0.872, *p-value*=0.3991). In contrast, the relative expression of PR-1a was higher in intact plants than in detached leaves, but only in MM plants (Table S4; plant treatment x plant variety interaction: χ2=14.0835, p-value<0.001; CM plants: Fig.2c; *t-ratio*=1.704, *p-value*=0.1121; MM plants: *t-ratio*=-4.090, *p-value*=0.0026). Importantly, in all cases, both intact plants and detached leaves mounted a significant response (Table S5).

**Fig. 2.**
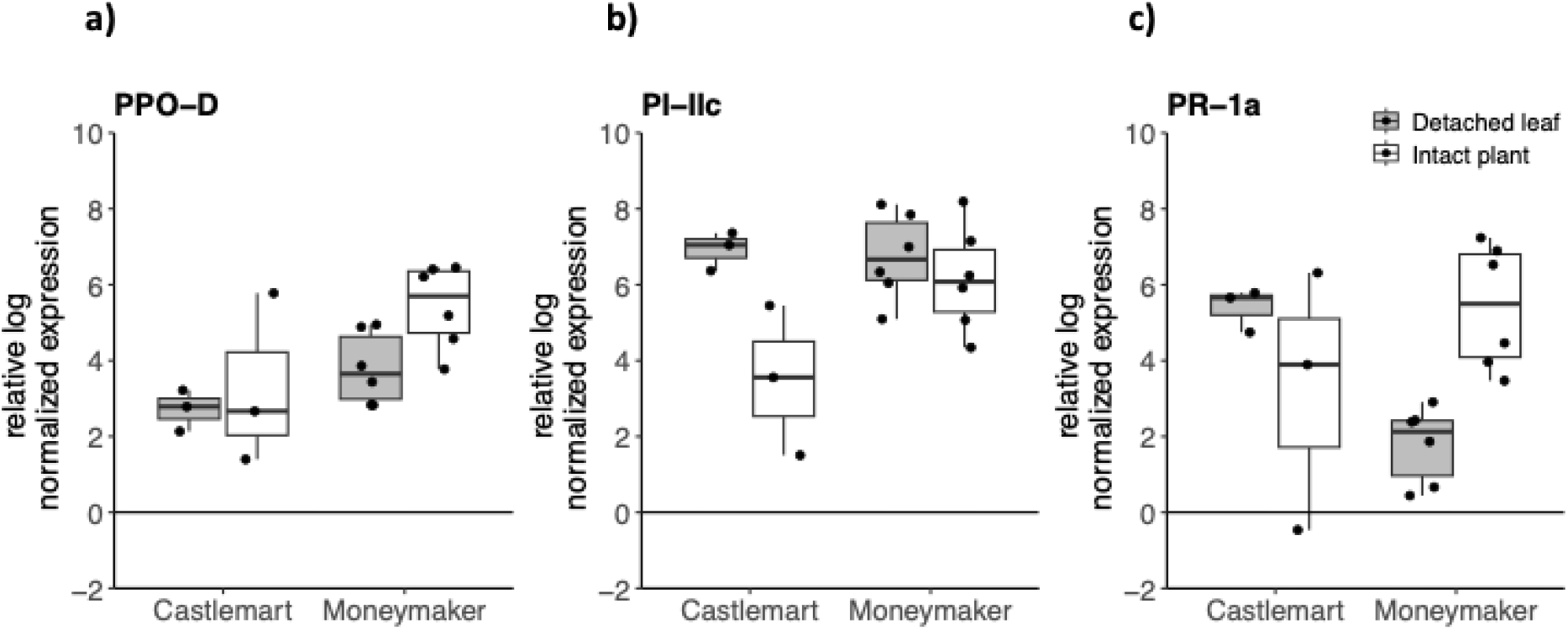
Relative log normalized gene expression of a) PPO-D, b) PI-IIc and c) PR-1a genes in detached leaves (grey) and intact plants (white) of CM and MM varieties, infested with *T. urticae*. The plotted values correspond to the difference to the control treatment (i-e. uninfested plants) and the zero line indicates no difference to the control. Each dot represents the average of three technical replicates per biological replicate.

### Induction of plant defences persists after detachment and increases with longer infestations

All three plant defence-related genes had higher expression in infested than in uninfested detached leaves (Fig.3a, *z-value*=2.791, *p-value*=0.005; Fig.3b, *z-value*=3.124, *p-value*=0.002; Fig.3c, *z-value*=4.161, *p-value*<0.001). Furthermore, an increase in defence induction was observed across time of infestation in detached leaves, in a gene specific manner. Indeed, while the expression of the early-gene PPO-D registered a significant but slight increase throughout the fourteen days of leaves infestation (Fig.3a, *z-value*=2.071, *p-value*=0.038), the expression of the two late-genes PI-IIc and PR-1a increased sharply over the course of the experiment (Fig.3b, *z-value*=5.438, *p-value*<0.001 and Fig.3c, *z-value*=4.605, *p-value*<0.001, respectively), reaching their peak expression on the fourteenth day (ca. 1.7, 5.5, 3.3 increased relative expression in PPO-D, PI-IIc and PR-1a, respectively, Fig. 3).

**Fig. 3.**
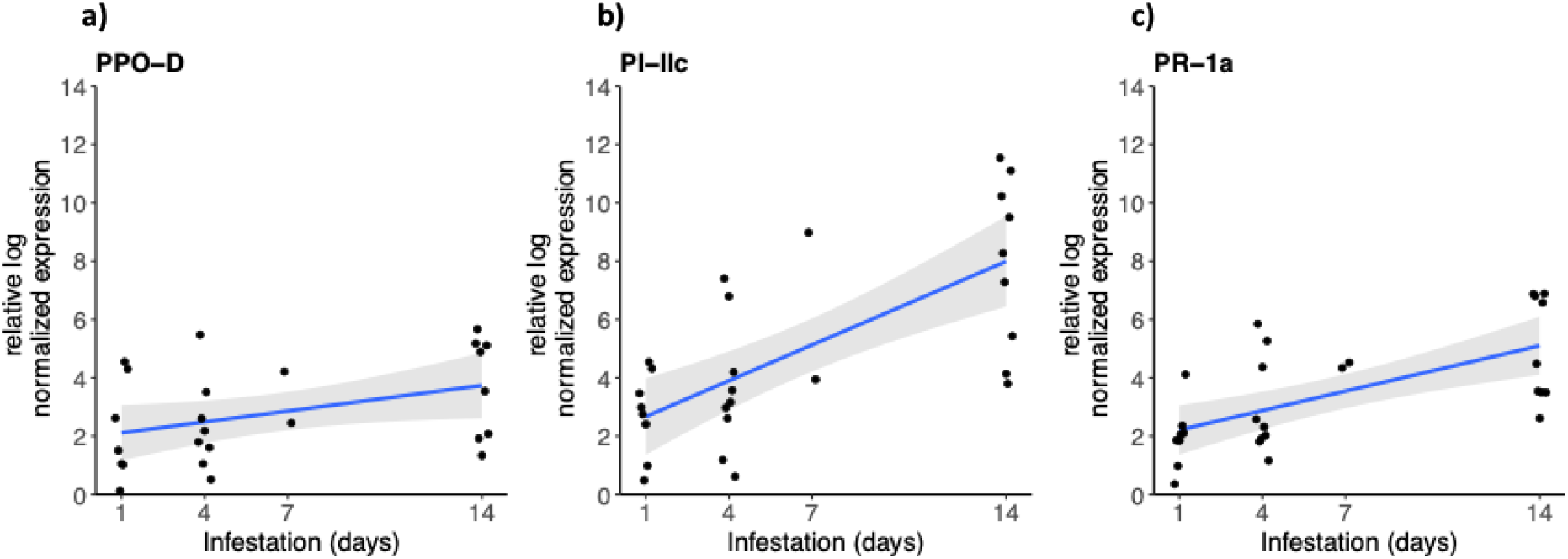
Relative log normalized gene expression of a) PPO-D, b) PI-IIc and c) PR-1a genes in detached MM leaves, infested with *T. urticae* for one, four, seven and 14 days. The plotted values correspond to the difference to the control treatment (i-e. uninfested plants) and the zero line indicates no difference to the control.Each dot represents the average of three technical replicates per replicate. The blue line represents the regression line with the shaded grey area representing the 95% confidence interval.

## Discussion

In this study we found that inducible defences are mounted in both intact plants and detached leaves. However, the magnitude of the early induction depended on the plant variety and on the gene being assessed. Indeed, the relative expression of PPO-D and of PR-1a varied between intact plants and detached leaves only in MM plants, being higher in intact plants, while the expression of PI-IIc significantly varied with plant integrity in CM plants only, being lower in intact plants. Importantly, we found that detached leaves continued to mount a defensive response locally after fourteen days of infestation, with all three genes increasing their expression with time.

Intact plants are generally thought to be preferable to evaluate the plant response to herbivory (Agrawal 2000a; Agrawal 2000b; Klingler et al. 2005; Michel et al. 2010). However, this is often unfeasible, and assays are performed on detached leaves instead (Dicke and Dijkman 1992; Stout et al. 1994; Koch et al. 1999; Musser et al. 2002; Sarmento et al. 2011a). The fact that *T. urticae* induced more defences in intact MM plants than in detached leaves in the beginning of the infestation period supports previous studies on plant-pathogen interactions which found lower expression of plant defence genes in detached leaves comparing to intact plants (Liu et al. 2007; Orłowska et al. 2012; Orłowska et al. 2013), thus confirming the importance of plant integrity in the induction of plant defences. However, the increased expression in intact plants compared to detached leaves was not consistent across all genes, with PI-IIc showing similar levels of expression in both plant treatments for MM variety. In addition, in CM plants, the pattern was reversed, with detached leaves mounting an equally high, or higher, local response than intact plants. These results align with those of previous studies on induction of indirect plant defences in herbivore-plant interactions (Arimura et al. 2001; Schmelz et al. 2001), which show that both intact plants and detached leaves mount a defence response but differently, with plant integrity influencing both the gene expression (Arimura et al. 2001) and the magnitude of volatile production (Arimura et al. 2001; Schmelz et al. 2001). All in all, these results suggest that the mechanisms of plant induction are intricately influenced by both plant integrity and variety.

Previous work with spider mite-induced direct plant defences has shown that herbivore performance is equally affected by defences induced on leaf discs or intact plants (Dias et al. 2022). Therefore, the multiplicity of patterns observed at the gene expression level we found may well cancel out, leading to an overall similar response on intact plants and detached leaves. Still, our results call for caution if the goal of the study is to address the role of a particular gene or the response in a particular plant variety.

The results reported here pertain to the expression of infested plants relatively to the expression of non-infested but still excised leaves. This means that any detected difference in gene expression must be due to herbivory and not to wounding due to excision. Furthermore, the response to mechanical wounding is considered negligible compared to the damage inflicted by herbivores (Korth and Dixon 1997; Reymond et al. 2000). Still, we cannot rule out the possibility that mechanical wounding has a priming effect on defence induction, resulting in increased expression of some genes following herbivore infestation. This may be the case of PI-IIc, whose expression has been shown to increase in mechanically wounded leaves (Howe et al. 1996).

While we have found that early induction of plant defences occurs in both intact plants and detached leaves, it could be that leaf excision compromised the persistence and magnitude of plant induced responses in the long-term, due to the increased senescence process known to occur (Smart 1994), or because the response ultimately depends on metabolites produced in other parts of the plant (Harborne 2007), or nutrients and resources acquired in other parts of the plant (Schultz et al. 2013). However, we found that none of the genes measured decreased expression with time, instead observing an increase of all genes after 14 days of infestation on detached leaves. Importantly, our observations that *T. urticae* induce a large increase in expression of so called “late” genes are in line with previous study performed on intact plants (Alba et al. 2015). This suggests that plant defence mechanisms are active for at least two weeks after detachment, which validates the use of detached leaves in experimental evolution studies that require the persistence of such defences (e.g., Fellous et al. 2014; Godinho et al. 2024).

In conclusion, we show that detached tomato leaves mount and maintain a response to herbivory through time, with the mechanisms of plant induction being intricately influenced by plant integrity and plant variety. Although we acknowledge the study of, and on, intact plants as being more ecologically relevant (Bezemer and Van Dam 2005; Van Dam and Heil 2011), our results demonstrate that the use of detached leaves in plant-herbivore interaction studies may also be a valid approach, depending on the study aims. Still, the broad diversity of herbivore feeding adaptations and respective counter-defences to plant defences calls for future studies addressing the importance of plant integrity in plant interactions with other herbivore and plant species.

## Author contributions

SM and LRR conceptualized the study and designed the experiments with support from DG. MK, DG and EY performed the experiments. MK performed statistical analyses with support from IF and LRR. MK wrote the first draft of the manuscript, with substantial help from LRR. All authors contributed to revisions.

## Acknowledgment

The authors are grateful to all Mite Squad members for helpful discussions and technical support in the laboratory, namely Pauline Lecroix.

## Data availability

The datasets generated during and/or analysed during the current study are at 10.6084/m9.figshare.28750130.

