## Supplementary Information for "Inducible tomato defences persist in detached leaves, despite differential plant variety and gene-dependent expression"

**Table S1** Number of replicates per treatment used to analyse the effect of plant integrity and persistence of induction on gene expression

**Table S2** Information about the genes, and corresponding primers, used in the quantification of plant defence induction

**Table S3** Description of the statistical models used to analyse the effect of plant integrity and defence induction persistence on gene expression

**Table S4** Statistical results of the effect of plant integrity and persistence of defence induction on gene expression

**Table S5** *A posteriori* contrasts between *T. urticae* infested and uninfested treatments to estimate the effect of infestation on gene expression

**Table S1** Number of replicates per treatment used to analyse the effect of plant integrity and persistence of induction on gene expression. a) refers to experiments performed to compare plant defence induction in intact plants and detached leaves and b) to experiments testing the persistence of induction in detached leaves. "Plant variety": tomato plant varieties used in the experiment (Castlemart or Moneymaker). "Treatment": degree of plant integrity (intact plants or detached leaves). "Infestation": degree of plant infestation (uninfested plants or plants infested with *T. urticae*); "No. of replicates": number of samples analysed for each set of independent variables (note that three technical replicates were run per sample). Samples in which the reactions failed to reach the minimum signal intensity at 40 cycles of qPCR were excluded from the analysis ([a], out of 3 samples, none reached this minimum); "Days of treatment": number of days during which tomato leaves were infested with *T. urticae* or kept uninfested before being sampled (one, four, seven and 14 days)

**a)**

| Defence-related genes | Plant variety | Treatment | Infestation | No. of replicates |
|---|---|---|---|---|
| PPO-D | Castlemart | Intact | Uninfested | 3 |
| | Castlemart | Intact | Infested | 3 |
| | Castlemart | Detached | Unifested | 3 |
| | Castlemart | Detached | Infested | 3 |
| | Moneymaker | Intact | Unifested | 6 |
| | Moneymaker | Intact | Infested | 6 |
| | Moneymaker | Detached | Unifested | 6 |
| | Moneymaker | Detached | Infested | 6 |
| PI-IIc | Castlemart | Intact | Unifested | 0[a] |
| | Castlemart | Intact | Infested | 3 |
| | Castlemart | Detached | Unifested | 3 |
| | Castlemart | Detached | Infested | 3 |
| | Moneymaker | Intact | Unifested | 4 |
| | Moneymaker | Intact | Infested | 6 |
| | Moneymaker | Detached | Unifested | 6 |
| | Moneymaker | Detached | Infested | 6 |
| PR-1a | Castlemart | Intact | Unifested | 3 |
| | Castlemart | Intact | Infested | 3 |
| | Castlemart | Detached | Unifested | 3 |
| | Castlemart | Detached | Infested | 3 |
| | Moneymaker | Intact | Unifested | 6 |
| | Moneymaker | Intact | Infested | 6 |
| | Moneymaker | Detached | Unifested | 6 |
| | Moneymaker | Detached | Infested | 6 |

**b)**

| Defence-related genes | Days of treatment | Infestation | No. of replicates |
|---|---|---|---|
| PPO-D | 1 | Uninfested | 8 |
| | | Infested | 9 |
| | 4 | Uninfested | 9 |
| | | Infested | 9 |
| | 7 | Uninfested | 9 |
| | | Infested | 2 |
| | 14 | Uninfested | 9 |
| | | Infested | 9 |
| PI-IIc | 1 | Uninfested | 5 |
| | | Infested | 9 |
| | 4 | Uninfested | 9 |
| | | Infested | 9 |
| | 7 | Uninfested | 9 |
| | | Infested | 2 |
| | 14 | Uninfested | 6 |
| | | Infested | 9 |
| PR-1a | 1 | Uninfested | 9 |
| | | Infested | 9 |
| | 4 | Uninfested | 9 |
| | | Infested | 9 |
| | 7 | Uninfested | 9 |
| | | Infested | 2 |
| | 14 | Uninfested | 9 |
| | | Infested | 9 |

**Table S2** Information about the genes, and corresponding primers, used in the quantification of plant defence induction

| Defence-related genes | Name | Gene identifier | Forward primer (5'→3') | Reverse primer (5'→3') | References |
|---|---|---|---|---|---|
| **Actin** | Actin | Solyc03g078400.2 | TCAGCACATTCCAGCAGATGT | AACAGACAGGACACTCGCACT | (Tomato Genome Consortium 2012) |
| **PPO-D** | Polyphenol-oxidase-D | Solyc08g074680.2 | GCCCAATGGAGCCATATC | ACATTCGATCCACATTGCTG | (Newman et al. 1993) |
| **PI-IIc** | Proteinase Inhibitor IIc | Solyc03g020050.2 | CAGGATGTACGACGTGTTGC | GAGTTTGCAACCCTCTCCTG | (Gadea et al. 1996) |
| **PR-1a** | Pathogenesis-related protein 1a | Solyc09g007010.1 | TGGTGGTTCATTTCTTGCAACTAC | ATCAATCCGATCCACTTATCATTTTA | (Van Kan et al. 1992) |

**Table S3** Description of the statistical models used to analyse the effect of plant integrity and defence induction persistence on gene expression. a) refers to experiment performed to assess plant defence induction comparing intact plants and detached leaves and b) to experiment testing the persistence of induction in detached leaves. "Sample size" refers to the total number of replicates included in each analysis. "Maximal model" describes the complete set of explanatory variables included in each model, while "minimal model" excludes non-significant random and fixed factors. Random factors are indicated within brackets. "Treatment": degree of plant integrity (intact plants and detached leaves); "Plant variety": tomato plant varieties used in the experiment (Castlemart and Moneymaker); "Block": the day at which the replicates were tested (blocks 1 and 2); "Days of treatment": the number of days tomato detached leaves were infested with *T. urticae* or kept uninfested before being sampled (one, four, seven and 14 days); "Leaf": the tomato leaf tested (third, fourth and fifth detached leaves, counting from the roots up to the tip of the plant)

**a)**

| Defence-related genes | Sample size | Maximal model | Minimal model | Error distribution |
|---|---|---|---|---|
| PPO-D | 36 | Treatment*Plant variety + (1\|Block:Plant variety) | Treatment+Plant variety | Normal |
| PI-IIc | 31 | Treatment*Plant variety + (1\|Block:Plant variety) | Treatment*Plant variety | Normal |
| PR-1a | 36 | Treatment*Plant variety + (1\|Block:Plant variety) | Treatment*Plant variety | Normal |

**b)**

| Defence-related genes | Sample size | Maximal model | Minimal model | Error distribution |
|---|---|---|---|---|
| PPO-D | 64 | Days of treatment + (1\|Leaf) | Days of treatment | Normal |
| PI-IIc | 58 | Days of treatment + (1\|Leaf) | Days of treatment | Normal |
| PR-1a | 65 | Days of treatment + (1\|Leaf) | Days of treatment | Normal |

**Table S4** Statistical results of the effect of plant integrity and persistence of defence induction on gene expression. This table refers to the experiment performed to assess plant defence induction comparing intact plants with detached leaves. "df": degrees of freedom. "$\chi^2$": chi-square value obtained in each analysis. Statistically significant differences ($p \leq 0.05$) are represented in bold

| Defence-related genes | Explanatory variables | df | $\chi^2$ | P-value |
|---|---|---|---|---|
| PPO-D | **Treatment** | **1** | **5.8941** | **0.0152** |
| | **Plant variety** | **1** | **8.3925** | **0.0038** |
| | Treatment*Plant variety | 1 | 0.9585 | 0.3276 |
| PI-IIc | **Treatment** | **1** | **12.9000** | **<0.001** |
| | Plant variety | 1 | 0.0506 | 0.8220 |
| | **Treatment*Plant variety** | **1** | **5.9013** | **0.0151** |
| PR-1a | Treatment | 1 | 2.9036 | 0.0884 |
| | **Plant variety** | **1** | **10.9632** | **<0.001** |
| | **Treatment*Plant variety** | **1** | **14.0835** | **<0.001** |

**Table S5** Comparisons between *T. urticae* infested and uninfested treatments to estimate the effect of infestation on gene expression. This table refers to the experiment performed to assess plant defence induction comparing intact plants with detached leaves. Comparisons between treatments (intact plant or detached leaves) and plant varieties (Castlemart or Moneymaker) were performed using the *emmeans* and the *test* function of the *emmeans* package. "Lower CL – Upper CL": 95% confidence limits for normalized gene expression of *T. urticae* infested treatment. "T ratio": the T-test value obtained in each comparison. Statistically significant differences ($p \leq 0.05$) are represented in bold

| Defence-related genes | Plant variety | Treatment | Lower CL – Upper CL | T ratio | P-value |
|---|---|---|---|---|---|
| PPO-D | **CM** | **Intact** | **2.51 – 4.77** | **6.896** | **<0.001** |
| | **CM** | **Detached** | **1.23 – 3.49** | **4.468** | **<0.001** |
| | **MM** | **Intact** | **4.36 – 6.15** | **12.610** | **<0.001** |
| | **MM** | **Detached** | **3.08 – 4.87** | **9.539** | **<0.001** |
| PI-IIc | **CM** | **Intact** | **2.05 – 4.96** | **5.205** | **<0.001** |
| | **CM** | **Detached** | **5.47 – 8.38** | **10.285** | **<0.001** |
| | **MM** | **Intact** | **5.12 – 7.18** | **12.922** | **<0.001** |
| | **MM** | **Detached** | **5.71 – 7.77** | **14.155** | **<0.001** |
| PR-1a | **CM** | **Intact** | **1.32 – 5.17** | **3.647** | **0.0030** |
| | **CM** | **Detached** | **3.47 – 7.32** | **6.057** | **<0.001** |
| | **MM** | **Intact** | **4.01 – 6.79** | **8.615** | **<0.001** |
| | **MM** | **Detached** | **0.42 – 7.32** | **2.831** | **0.0142** |
